# A defined diet for pre-adult *Drosophila melanogaster*

**DOI:** 10.1101/2023.08.28.555220

**Authors:** Felipe Martelli, Annelise Quig, Sarah Mele, Jiayi Lin, Tahlia L. Fulton, Mia Wansbrough, Christopher K. Barlow, Ralf B. Schittenhelm, Travis K. Johnson, Matthew D.W. Piper

## Abstract

*Drosophila* melanogaster is unique among animal models because it has a fully defined synthetic diet available to study nutrient-gene interactions. However, use of this diet is limited to adult studies due to impaired larval development and survival. Here, we provide an adjusted formula that improves larval growth, development rate, and rescues survival. We demonstrate its use for exploring pre-adult diet compositions of therapeutic potential in a model of an inherited metabolic disorder.

## Main

Nutrition is a major environmental factor that affects animal physiology, metabolism, and influences traits such as survival and fitness^1^. Understanding how each nutrient contributes to these phenotype remains a major challenge in biology due to the complex composition of food. The vinegar fly, *Drosophila melanogaster*, is unique amongst animal models as it has a fully synthetic chemically-defined (holidic) medium allowing for manipulation of individual nutrients^2^. This diet, named hereon as 100N (“N” describes optimal nitrogen content for adults), has been used extensively to test how nutrients impact complex traits such as reproduction, lifespan, behaviour, and host-microbiome interactions^2–5^. However, a major limitation of 100N is that flies reared solely on it (i) are significantly delayed when compared to flies reared on sugar-yeast (SY) diet (100N: 17.95 days to eclosion; SY: 10.55 days at 25ºC), (ii) show reduced survivorship (100N: 85%; SY: 96%), and (iii) are smaller in size upon eclosion (**Fig. 1a**), suggesting that 100N is suboptimal for growth and development.

**Figure 1.**
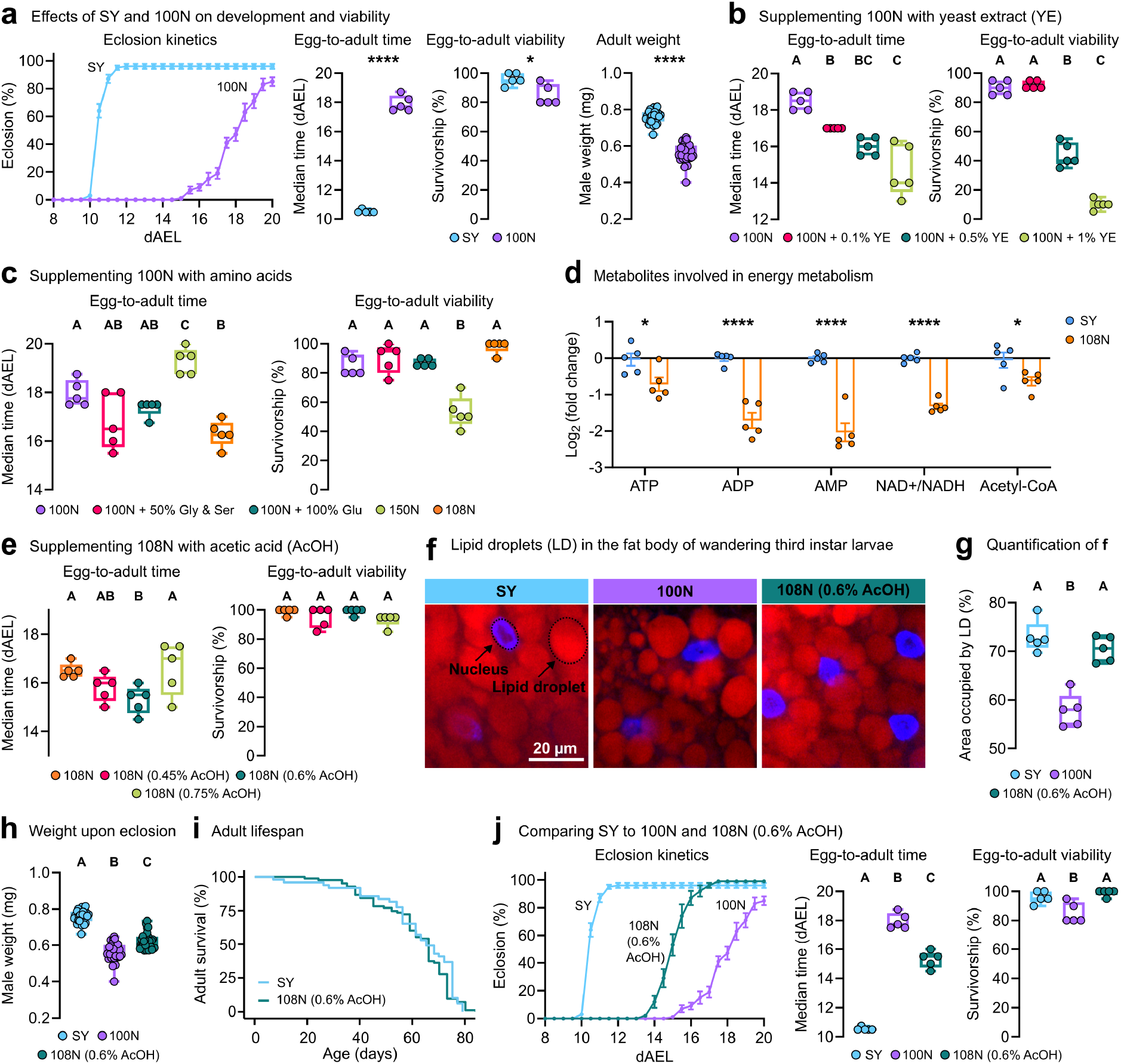
Optimizing the holidic diet for *Drosophila* pre-adulthood. **a**, Eclosion kinetics, median egg-to-adult duration in days after-egg-lay (dAEL), mean percentage egg-to-adult survivorship, and male weight upon eclosure (mg) on SY and 100N diets. **b-c**, Egg-to-adult duration (dAEL) and egg-to-adult survivorship on 100N supplemented with (**b**) 0.1%; 0.5%; or 1% of yeast extract (YE), and (**c**) 50% Ser and Gly; 100% Glu; 50% supplementation of all 20 aa (150N); or 8% supplementation of all 20 aa (108N). **d**, Log_2_(fold change) of energy related metabolites in larvae significantly affected in 108N in comparison to SY media. **e**, Egg-to-adult duration (dAEL) and egg-to-adult survivorship on 108N containing 0.45%; 0.6%; or 0.75% acetic acid (AcOH, v/v). **f**, Lipid storage in the late-larval fat body on 100N, 108N (0.6% AcOH), or SY. Nucleic acids (blue) and lipid droplets (red), 400 x magnification. **g**, Percentage of fat body area occupied by lipid droplets. **h**, Adult male weight upon eclosion (mg) raised on SY, 100N, or 108N (0.6% AcOH). **i**, Lifespan curves of adult flies fed SY food after being raised on either 108N (0.6% AcOH) or SY during their pre-adult period. **j**, Eclosion kinetics, egg-to-adult duration (dAEL) and egg-to-adult survivorship on SY, 100N, and 108N (0.6% AcOH) diets. **a, b, c, e**, and **f**, five biological replicates per group of 20 individuals each. **d**, five biological replicates per group of 10 larvae each. **h**, five larvae per group (each point represents an average of 5 individual measurements per larva). **i**, 5 to 10 replicates per group of 10 female adults each. **j**, five biological replicates per group of at least 5 adults each. Data shown in **a** are repeated in **j**, to compare against 108N (0.6% AcOH). **b, c, e, f, h**, and **j**, One-way ANOVA followed by Tukey’s HSD test. **i**, Cox Proportional-Hazards modelling (p = 0.472). Different letters represent statistically significant differences (p < 0.05). **a** and **d**, Student’s unpaired *t*-test, ****p < 0.0001 ***p < 0.001, **p < 0.01, *p < 0.05.

To improve diet performance, we made a series of adjustments to the 100N diet. Before checking individual nutrients, we started by investigating the effects of adding yeast extract (YE). Adding 0.1% water soluble YE improved development time by 1.5-days, and while further additions progressively improved development they reduced survival indicating nutritional imbalance (**Fig. 1b, Supplementary Figure 1a**). Since YE is a rich source of amino acids (aa) and aa signal growth, we asked whether aa supplementation alone could provide similar benefits without compromising survival. Adding 50% more (in comparison to 100N) Serine (Ser) and Glycine (Gly) to supplement nitrogen and one-carbon units for anabolism, or 100% more Glutamate (Glu) to supplement nitrogen and carbon, appeared to reduce development time without compromising viability (**Fig. 1c, Supplementary Figure 1b-d**). Addition of 8% more of all amino acids (108N; equal to the nitrogen content of the individual aa supplementations) reduced egg-to-adult development time by 1.7-days and significantly increased survivorship (98%, **Fig. 1c**). Further supplementation of all aa (150N) was highly detrimental to survival and development rate (**Fig. 1c**).

To improve the formula further, we investigated the differences between 108N- and SY-raised *Drosophila* larvae via untargeted metabolomics (**Supplementary Table 1-2**). Metabolic profiles suggested reduced energy metabolism (**Fig. 1d**), as well as lipid-, carnitine-, and β-oxidation-associated metabolites (**Supplementary Figure 2**) in the 108N-raised larvae. Various supplementation trials (carnitine, lipids, sucrose) failed to improve larval development on 108N (**Supplementary Figure 3a-h**). Since the natural substrate of *Drosophila* larvae is fermenting fruit rich in alcohol and acetate, but poor in long-chain fatty acids^6,7^, we tested acetic acid (AcOH) supplementation. Doubling the AcOH (to 0.6% v/v) significantly reduced egg-to-adult time by 1-day compared to 108N, without compromise to survival (**Fig. 1e, Supplementary Figure 3i**). Tissue analysis of animals reared on 108N (0.6% AcOH) showed larger adipose tissue (fat body) lipid stores in comparison to 100N-raised larvae, reaching equivalent levels to SY-raised larvae, indicating a general metabolic improvement (**Fig. 1f-g**). These individuals also attained significantly higher bodyweights as newly-eclosed adults, compared to 100N-raised animals, despite reaching adulthood in a shorter time (**Fig. 1h**). Importantly, the lifespan of flies raised on 108N (0.6% AcOH) was no different to those raised on SY (**Fig. 1i**). Overall, when compared to the original holidic diet, our adjusted formula improved survivorship to 98% (equivalent to SY-raised *Drosophila*), reduced egg-to-adult time by 2.7 days (**Fig. 1j**), restored fat levels, and enhanced body mass, without compromising adult lifespan.

A synthetic diet on which *Drosophila* can be raised is particularly valuable in the modelling of human metabolic disorders, where phenotypic abnormalities often manifest prior to adulthood^8^. For example, human disorders such as Phenylketonuria manifest with rapid neurological decline prior to adulthood mitigated by a custom diet with single aa restriction that drastically improves health outcomes^9^. However, the majority of metabolic disorders lack therapeutic diets^8^. To demonstrate the potential of our diet as a tool to find new dietary treatments, we applied it in a model of human Branched Chain Amino Acid (BCAA) Transaminase (*BCAT2*) deficiency, a rare disorder associated with developmental delay, intellectual disability, seizures, and movement defects^10^. *BCAT2* catalyzes the first step in Valine (Val), Leucine (Leu) and Isoleucine (Ile) catabolism^11^ and patients have high plasma and tissue levels of these aa^10^ (**Fig. 2a**).

**Figure 2.**
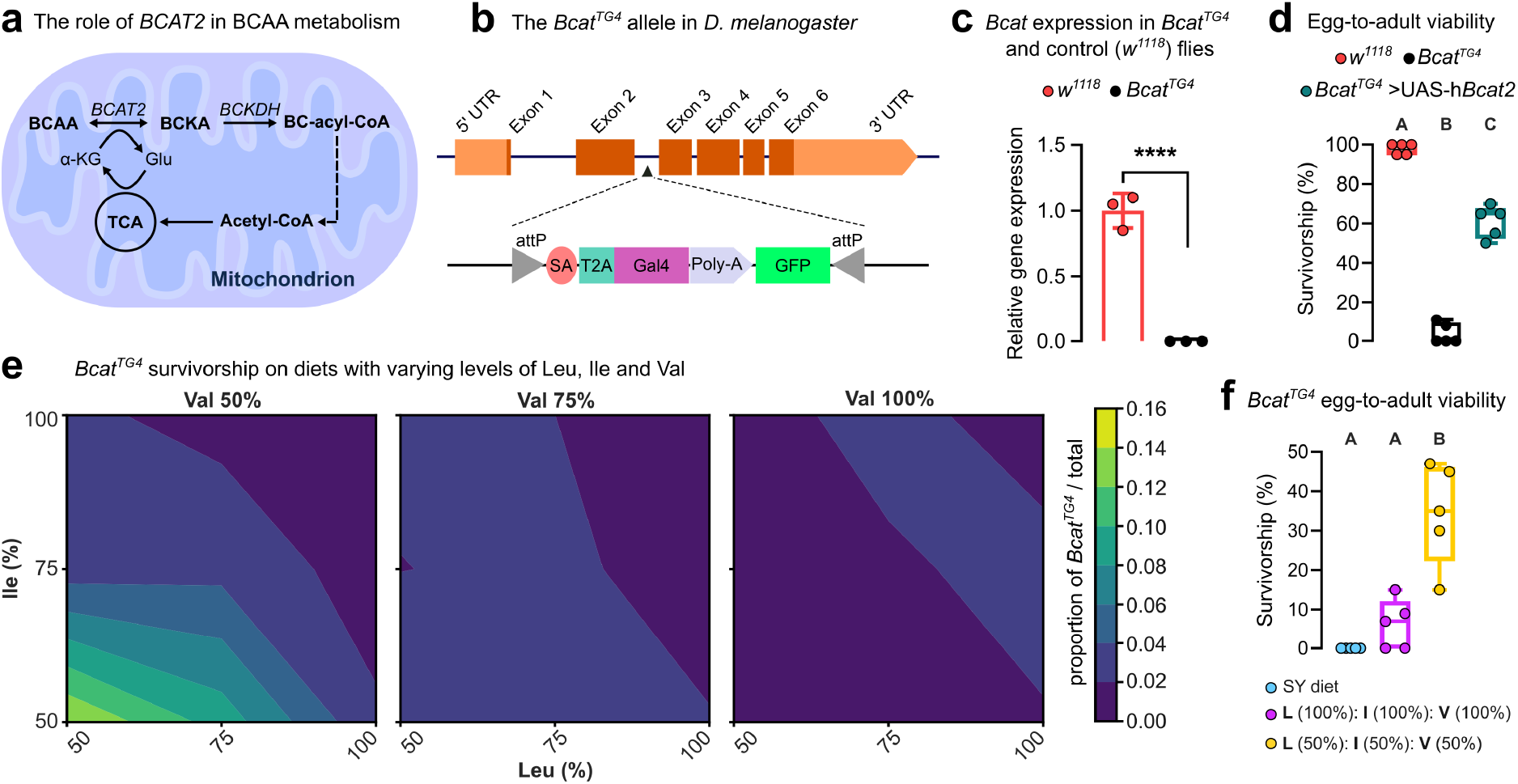
Applying the pre-adult-optimized synthetic diet in a *Drosophila* inherited metabolic disorder model. **a**, Schematic of branched-chain amino acid (BCAA) metabolism. BCAA are converted into branched-chain α-keto acid (BCKA) by the action of BCAT2, the deficiency of which causes a rare metabolic disorder. **b**, Schematic of Bcat^TG4^ allele in *D. melanogaster* containing the disruption T2A-GAL4 (TG4) cassette. **c**, Validation of the *Bcat*^*TG4*^ allele, showing *Bcat* expression is not detected. *w*^*1118*^ are a wildtype control genotype. **d**, Egg-to-adult survivorship of *Bcat*^*TG4*^ flies and *Bcat*^*TG4*^ expressing the human orthologue *Bcat2*, when reared on 108N (0.6% AcOH). **e**, *Bcat*^*TG4*^ survivorship contour plots resulting from a 3-way nutrient (Leu, Ile, Val) dose response array (27 diets total, 8-9 replicates performed per diet). **f**, Egg-to-adult survivorship of *Bcat*^*TG4*^ flies raised on SY, 108N (0.6% AcOH), and 108N (0.6% AcOH) with 50% BCAA content. **d** and **f**, five biological replicates per group of 20 individuals each. **c**, Student’s unpaired *t*-test, ****p < 0.0001. **d** and **f**, One-way ANOVA followed by Tukey’s HSD test. Different letters represent statistically significant differences (p < 0.001).

We used a *Drosophila Bcat* allele (*Bcat*^*TG4*^) that comprises a T2A-GAL4 cassette inserted within the second intron of *Bcat*, which abolishes expression of the full-length transcript and truncates protein translation^12^ (**Fig. 2b-c**). Egg-to-adult survival for *Bcat*^*TG4*^ homozygotes on 108N (0.6% AcOH) is 4% and this was rescued to 60% via expression of a human ortholog *BCAT2* transgene suggesting fly and human genes are functionally equivalent (**Fig. 2d**). We next tested *Bcat*^*TG4*^ flies and their survival relative to sibling controls in a nutrigenomic array of 27 BCAA-modified diets where each BCAA (Leu, Ile, and Val) was co-varied across three concentrations (50%, 75% and 100%). *Bcat*^*TG4*^ flies had highest relative survival when reared on a diet where all three BCAAs were reduced by 50%, while no benefit was observed from restricting one or two BCAA at a time in any combination (**Fig. 2e**). Tracking *Bcat*^*TG4*^ flies developing in isolation on the 50% all BCAA diet (to determine absolute survival, rather than relative to genetic controls) revealed a striking improvement in survivorship (from ∼5% to 34%, **Fig. 2f**). These findings mirror diet responses observed in BCAT2 deficiency patients^10^ and observations from a mouse BCAT2 model^13^. This highlights the potential of our modified synthetic diet for nutrigenomic studies of fly disease models for which effective diet formulations are lacking. More broadly, our diet expands the study of nutrient-gene interactions to the entire life-cycle of *Drosophila*, thus permitting the exploration of even complex processes such as growth and development.

## Methods

### Fly stocks and base diet

The synthetic (holidic) diet optimizing experiments were performed on *w*^1118^ (Bloomington stock #3605). For experiments involving *Bcat* mutant flies, *Bcat*^*TG4*^ mutants (Bloomington stock #79238) were first outcrossed to *w*^1118^ for removal of y^1^ and then maintained over fluorescently marked balancer chromosome (Bloomington stock #35522). Rescue with human ortholog was performed by cross with UAS-h*BCAT2* (Bloomington stock #84835). Dietary manipulations were conducted using the synthetic diet^2,3^ (**Supplementary Table 3**). The sugar-yeast (SY) diet was prepared as per standard media (**Supplementary Table 3**). All stocks of *Drosophila melanogaster* were maintained at 18 °C on sugar-yeast medium under natural photoperiod conditions. For experiments, flies were reared in incubators at 25 °C and 45% humidity in specific diets as described below.

### Testing holidic diet improvements

The modifications tested with the holidic diet include manipulation of amino acid levels, supplementation with yeast extract (Bacto™ Yeast Extract, Thermo Fisher Scientific), sucrose (Sigma-Aldrich, S1888), organic virgin coconut oil (Macro Wholefoods Market), Linolenic acid (L2376-500MG, Merck), L-Carnitine hydrochloride (Merck, C0283-5G), and acetic acid (Fisher, A/0400/PB15). The solutions of yeast extract, sucrose, acetic acid, and carnitine were prepared in Milli-Q water. Appropriate volumes of these solutions, with the exception of Linolenic acid and coconut oil, were added to the solid food surface and left for 24 hours in a 25°C incubator to allow for complete diffusion. No more than 300 μL was added to each vial. For Linolenic acid and coconut oil, appropriate volumes were added to the holidic diet prior to setting and homogenized with help of a magnetic stirrer before dispensing into vials. Diets with varied amino acid concentrations were prepared by modifying the base holidic diet recipe (**Supplementary Table 3**).

### Survival and developmental timing

Embryos were collected from population cages containing approximately 200 females and 50 males (5 to 10 days old) laying for 6 hours on apple juice agar plates supplemented with yeast paste. Embryos were then washed with distilled water and transferred to a new plate not containing yeast paste and left for 24 hours to hatch. Twenty first instar larvae were transferred to each vial for the different diets. For each diet, at least 5 replicate vials were tested (n = 100 individuals). Vials were monitored every 12 hours for a period of 20 days and the numbers of pupae and adults were recorded. Data were visualized as the cumulative percentage of individuals at each life stage as a function of time (days) from egg lay. Data were analyzed using one-way ANOVA followed by Tukey’s HSD test (GraphPad Prism 9).

### Metabolic profiling

Five replicates of 10 five-day-old larvae from each genotype and diet condition were collected, washed in PBS, blotted dried, and weighed. Samples were then transferred to 1.5 mL safe lock microtubes tubes (Eppendorf), flash frozen in liquid nitrogen, and stored at -80 °C. Thawed larvae were homogenized using a disposable pestle in 20 μL of ice-cold extraction solvent consisting of 2:6:1 chloroform:methanol:water with 2 μM of (CHAPS, CAPS, PIPES and TRIS) acting as internal standards. Once ground additional solvent was added to a final ratio of 20 μL of extraction solvent per mg of larvae. Samples were vortexed for 30 s and then sonicated in an ice-water bath for 10 min and centrifuged at 4°C (22,000 ×g for 10 min). The supernatant was then transferred to a glass vial for LC-MS based metabolomic analysis. 20 μL of each extract were combined to make a pooled quality control (PQC) sample.

LC-MS was performed using a Dionex Ultimate 3000 UHPLC coupled to an QExactive Plus mass spectrometer (Thermo Scientific). Samples were analyzed by hydrophilic interaction liquid chromatography (HILIC) following a previously published method^14^. The chromatography utilized a ZIC-pHILIC column 5μm, 150 x 4.6 mm with a 20 x 2.1 mm ZIC-pHILIC guard column (both Merck Millipore, Australia) (25 °C). A gradient elution of 20 mM ammonium carbonate (A) and acetonitrile (B) (linear gradient time-%B: 0 min-80%, 15 min-50%, 18 min-5%, 21 min-5%, 24 min-80%, 32 min-80%) was utilized. Flow rate was maintained at 300 μL/min. Samples were stored in the autosampler (6°C) and 10 μL was injected for analysis. MS was performed at 70,000 resolution operating in rapid switching positive (4 kV) and negative (−3.5 kV) mode electrospray ionization (capillary temperature 300°C; sheath gas flow rate 50; auxiliary gas flow rate 20; sweep gas 0; probe temp 120°C). Samples were randomized and processed in a single batch with intermittent analysis of PQC samples to ensure reproducibility. For accurate metabolite identification, a standard library of ∼500 metabolites was analyzed before sample testing and accurate retention time for each standard was recorded. This standard library also forms the basis of a retention time prediction model used to provide putative identification of metabolites not contained within the standard library^15^.

### Metabolic profiling data analysis

Acquired LC-MS/MS data was processed in an untargeted fashion using open source software IDEOM, which initially used msConvert (ProteoWizard)^16^ to convert raw LC-MS files to mzXML format and XCMS to pick peaks to convert to .peakML files^17^. Mzmatch was subsequently used for sample alignment and filtering^18^. Metabolites were identified based on accurate mass (<2 ppm) and comparison of their retention time against that determined for compounds in the standard library or predicted on the basis of their physiochemical characteristics^15^. Only metabolites that were identified with a level of confidence equal to or greater than 6 in IDEOM were used for downstream functional and statistical analyses, using MetaboAnalyst 5.0^19^. Global metabolic variations due to genotype and diet were visualized using PCA. One-way ANOVA [false discovery rates (FDR) < 0.05] was used to identify significant changes in metabolite levels. A heatmap was created to visualize the 50 topmost affected metabolites (based on fold change).

### Assessment of lipid environment by confocal microscopy

The effects of SY, 100N and 108N supplemented with 0.6% AcOH on the lipid storage of wandering third instar larvae were assessed using fluorescence microscopy. Fat bodies were dissected in PBS (Ambion) and subjected to lipid staining with Nile Red N3013 Technical grade (Sigma-Aldrich) and nucleic acid staining with DAPI (Sigma-Aldrich). Tissues were fixed in 4% PFA (Electron Microscopy Science) for 20 min and stained with 0.5 μg/mL Nile Red and 10 μg/mL DAPI in PBS + 0.1% Triton X-100 for 15 min. After staining, samples were washed in PBS and slide were mounted in Vectashield (Vector Laboratories). Microscopies were performed in CV1000 Disc Confocal Microscope (Olympus Life Science) at 400 x magnification. Images were analyzed using ImageJ software. The percentage of area occupied by lipid droplets per fat body samples was averaged from three independent measurements of sections of 2500 μm^2^. A total of five biological replicates were assessed for each dietary condition, each replicate consisting of a single tissue from a single larva. Results were analyzed using One-way ANOVA followed by Tukey’s HSD test (GraphPad Prism 9).

### Male weight upon eclosure

The adult weight upon eclosure of male flies reared on SY, 100N, or 108N + 0.6% AcOH was assessed using a SE2 Ultramicrobalance (Sartorius, Goettingen, Germany). At least 30 flies were weighed upon eclosure. Results were analyzed using One-way ANOVA followed by Tukey’s HSD test (GraphPad Prism 9).

### Adult lifespan

The survival of flies reared from egg-to-adult on 108N (0.6% AcOH) was compared to those reared on the SY diet. Upon eclosion, female adults from either diet were transferred onto SY diet and survivorship was checked every other day, at the same time when flies were transferred to fresh media. Five to ten replicates per group of ten flies each were performed. Fly survival was recorded using the software DLife^20^ and Cox Proportional-Hazards modelling was used for analysis (R, package survival). Difference in survival between groups was analysed using a linear model (base R, “lm”) and post-hoc comparisons from the emmeans package.

### BCAA-modified diet array

Large numbers of developmentally synchronized embryos obtained from multiple cages containing Bcat/FM7a,sChFP flies were collected and washed in distilled water, and transferred to 15 mL tubes containing PBS. Once the embryos had settled (1 minute), excess PBS was removed. Using a micropipette set to 5 μL and a tip cut to widen the bore, small volumes of embryos were aspirated and then dispensed into vials onto the medium surface. This was performed for each of the 27 modified synthetic diets tested with varied levels of leucine, isoleucine and valine replicated at least 8 times (**Supplementary Table 4**). Vials were checked daily for 30 days and the total number of adult flies were counted with the survival proportion of *Bcat* (*Bcat* / total flies) calculated. The linear mixed model (R package: lme4) was fitted to analyse the effects of diet, genotype, and their interaction on mean *Bcat* survival proportion (number of *Bcat* flies out of total), with replicates treated as a random effect.

### Real time PCR

The expression of *Bcat* was measured in *Bcat*^*TG4*^ homozygous flies and *w*^*1118*^ control flies. A total of five whole flies were used for a single biological replicate and three replicates were taken from each genotype. RNA isolation with TriSure (Bioline) was performed following the manufacturer’s instructions. RNA purity and concentration were evaluated by spectrophotometry (NanoDrop® ND-1000, NanoDrop Technologies). RNA samples were then diluted to 1 μg/μL and stored at -80 °C. cDNA was generated using the Superscript III reverse transcriptase (NEB). RT-qPCR analysis was performed in a 7500 Real Time PCR System (Applied Biosystems) using the SYBR® Green Reagent protocol (Applied Biosystems) and the following pairs of primers: *Bcat* Forward 5’ CCGCCTATTCACCGATCACA3’; *Bcat* Reverse 5’ TACGCCTTCATGCCCTCAAA3’; *RpL11* 5’ CGATCCCTCCATCGGTATCT3’; *RpL11* Reverse 5’ AACCACTTCATGGCATCCTC3’. Relative expression was calculated using the 2^(-DeltaDeltaC(T)) method and *RpL11* expression as a housekeeping gene. The results were analyzed by the Shapiro-Wilk normality test (p>0.05) to test for normality, before comparing the transcript levels between the two treatment groups using two-tailed Student’s t-tests (p < 0.05 (GraphPad Prism 9).

