## Supplementary Figures 1-3 for "A defined diet for pre-adult *Drosophila melanogaster*"

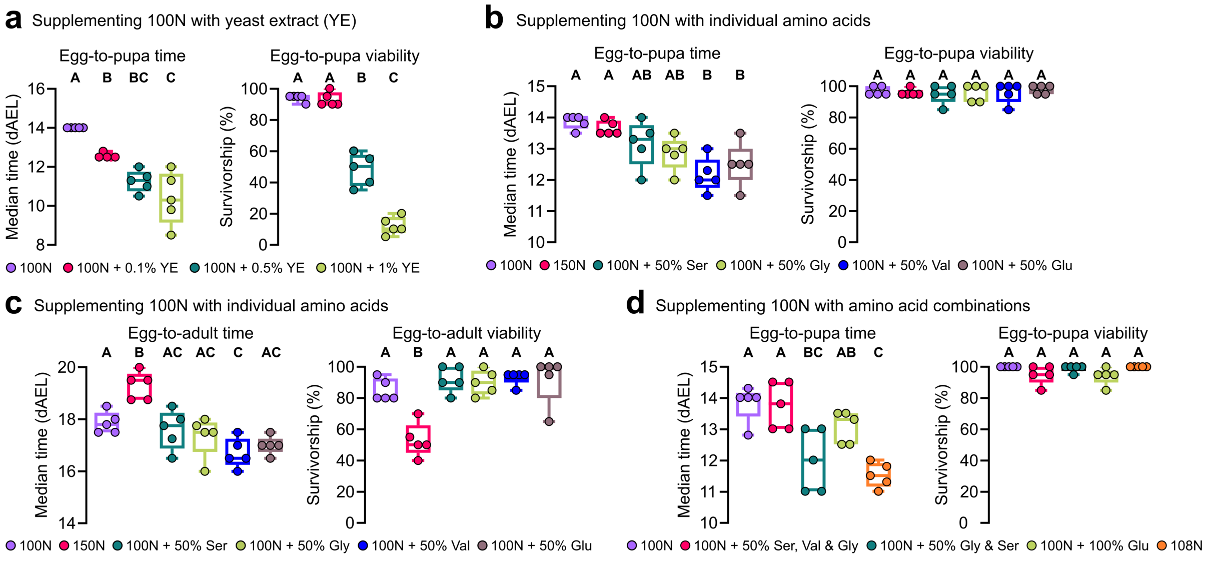


**Supplementary Figure 1. Optimizing the 100N diet for larval development. a, b, d,** Median egg-to-pupa duration in days after-egg-lay (dAEL) and mean percentage egg-to-pupa survivorship of flies reared on 100N supplemented with (**a**) 0.1%, 0.5%, or 1% of yeast extract (YE), (**b**) 50% Ser; 50% Gly; 50% Val; 50% Glu; or 50% supplementation of all 20 amino acids (150N), and (**d**) 50% Ser, Val and Gly; 50% Ser and Gly; 110% Glu; or 108N. **c,** Median egg-to-adult duration in days after-egg-lay (dAEL) and mean percentage egg-to-adult survivorship of flies reared on 100N supplemented with 50% Ser; 50% Gly; 50% Val; 50% Glu; or 50% supplementation of all 20 amino acids (150N). One-way ANOVA followed by Tukey's HSD test. Different letters represent statistically significant differences ($p < 0.05$).

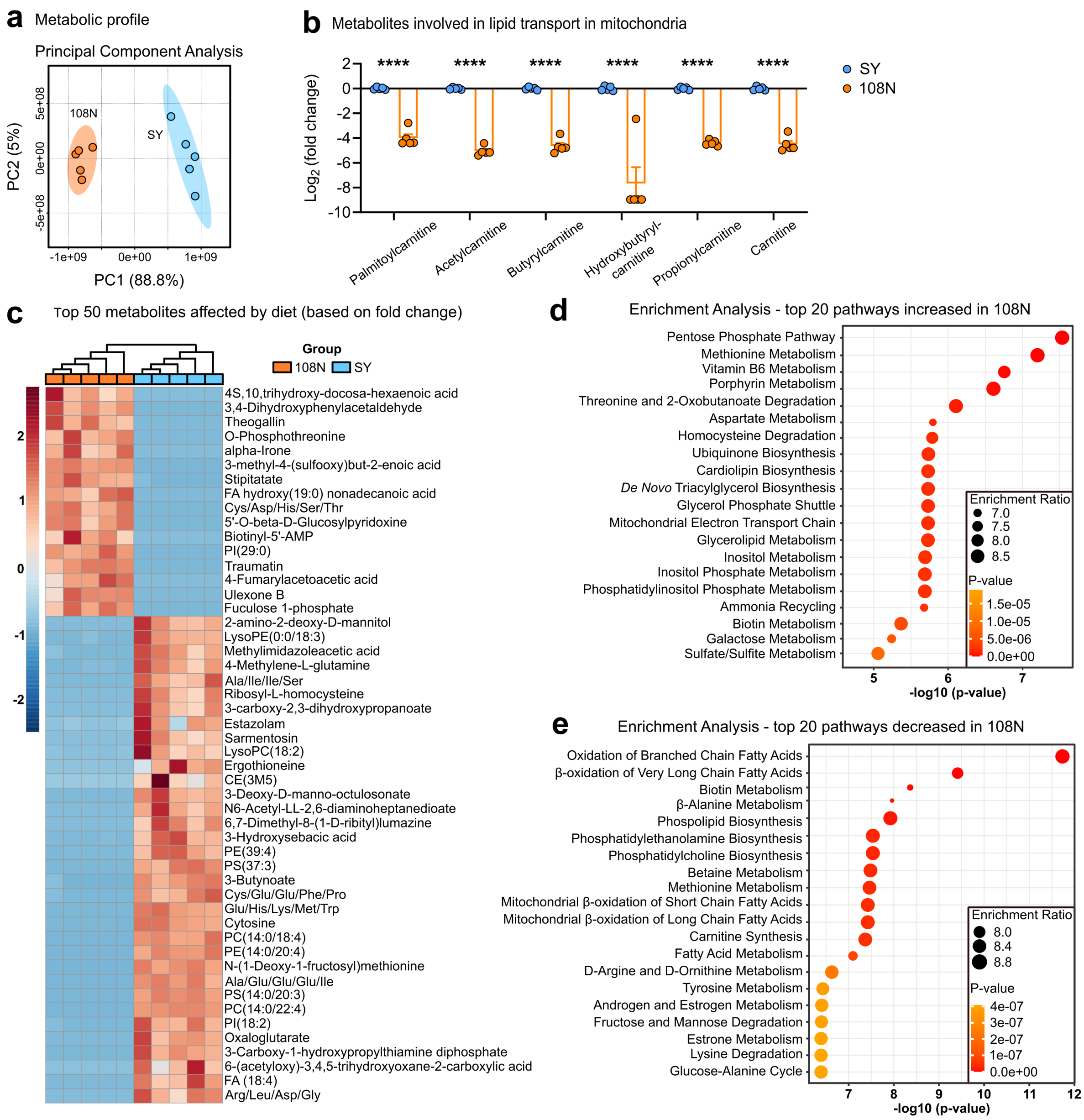


**Supplementary Figure 2. Metabolomics functional analysis of 3rd instar larvae reared on SY or 108N. a**, Principal Component Analysis (PCA) of metabolic profiles of third instar larvae reared on SY or 108N. Each dot represents the metabolome data sum of each sample. **b**, Log2(fold change) of larval metabolites involved in lipid transport in mitochondria, significantly affected in 108N in comparison to SY media. Student's unpaired *t*-test, ****p < 0.0001. **c**, Top 50 metabolites, based on foldchange, significantly affected by dietary treatment (Student's unpaired *t*-test, P < 0.05). The cell colors represent the z-scores, *i.e.*, the standardized scores on the same scale, calculated dividing a score's deviation by the standard deviation in the row. The features are color-coded by row with red indicating high intensity and blue indicating low intensity. **d**, Enrichment analysis showing the top 20 pathways (based on small molecule pathway database – SMPDB) increased in larvae reared on 108N in comparison to larvae reared on SY. **e**, Enrichment analysis showing the top 20 pathways (based on SMPDB) decreased in in larvae reared on 108N in comparison to larvae reared on SY.

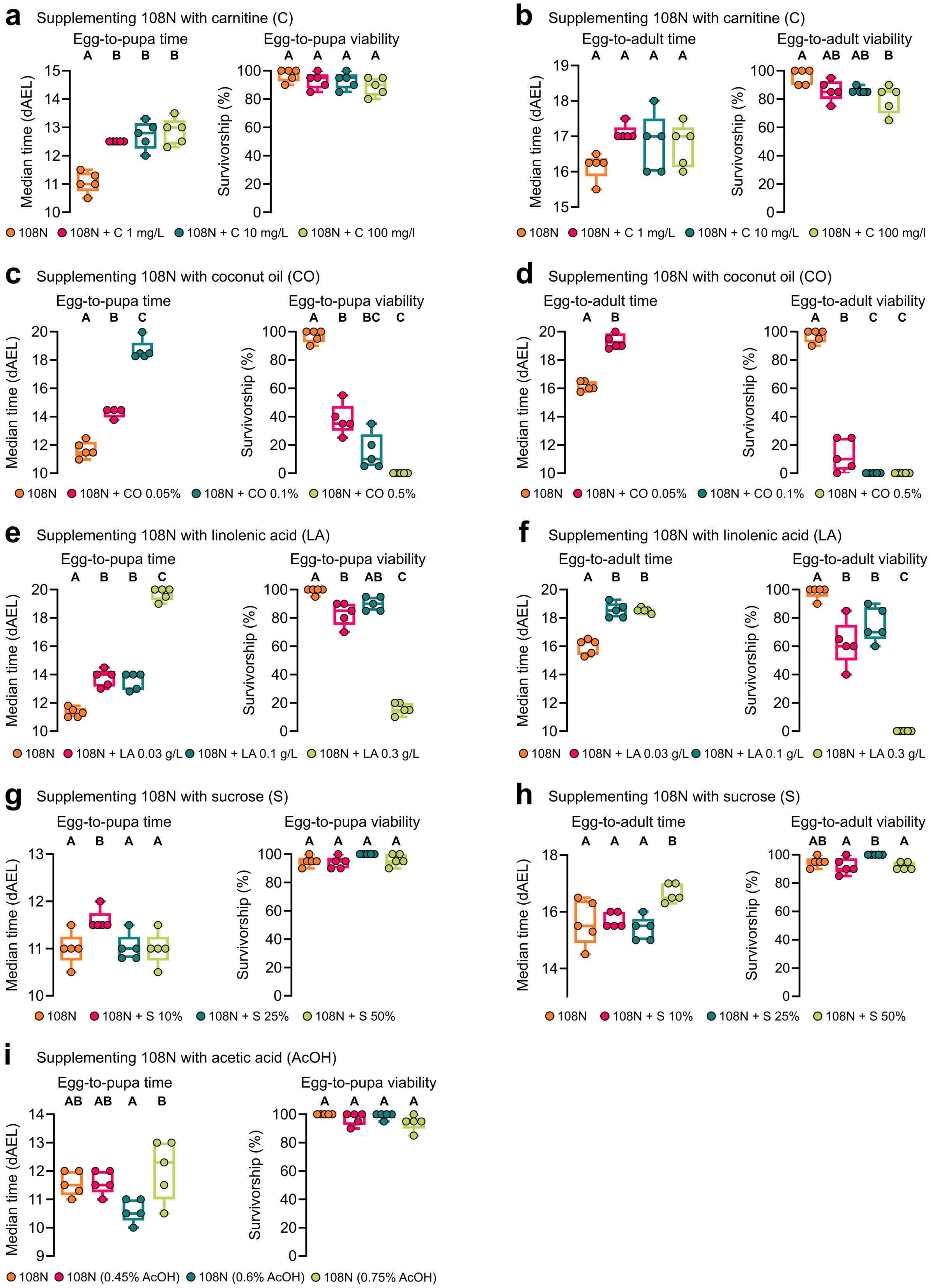
a
Supplementing 108N with carnitine (C)
Egg-to-pupa time
A B B B
Median time (dAEL)
Egg-to-pupa viability
A A A A
Survivorship (%)
108N
108N + C 1 mg/L
108N + C 10 mg/L
108N + C 100 mg/l
b
Supplementing 108N with carnitine (C)
Egg-to-adult time
A A A A
Median time (dAEL)
Egg-to-adult viability
A AB AB B
Survivorship (%)
108N
108N + C 1 mg/L
108N + C 10 mg/L
108N + C 100 mg/l
c
Supplementing 108N with coconut oil (CO)
Egg-to-pupa time
A B C
Median time (dAEL)
Egg-to-pupa viability
A B BC C
Survivorship (%)
108N
108N + CO 0.05%
108N + CO 0.1%
108N + CO 0.5%
d
Supplementing 108N with coconut oil (CO)
Egg-to-adult time
A B
Median time (dAEL)
Egg-to-adult viability
A B C C
Survivorship (%)
108N
108N + CO 0.05%
108N + CO 0.1%
108N + CO 0.5%
e
Supplementing 108N with linolenic acid (LA)
Egg-to-pupa time
A B B C
Median time (dAEL)
Egg-to-pupa viability
A B AB C
Survivorship (%)
108N
108N + LA 0.03 g/L
108N + LA 0.1 g/L
108N + LA 0.3 g/L
f
Supplementing 108N with linolenic acid (LA)
Egg-to-adult time
A B B
Median time (dAEL)
Egg-to-adult viability
A B B C
Survivorship (%)
108N
108N + LA 0.03 g/L
108N + LA 0.1 g/L
108N + LA 0.3 g/L
g
Supplementing 108N with sucrose (S)
Egg-to-pupa time
A B A A
Median time (dAEL)
Egg-to-pupa viability
A A A A
Survivorship (%)
108N
108N + S 10%
108N + S 25%
108N + S 50%
h
Supplementing 108N with sucrose (S)
Egg-to-adult time
A A A B
Median time (dAEL)
Egg-to-adult viability
AB A B A
Survivorship (%)
108N
108N + S 10%
108N + S 25%
108N + S 50%
i
Supplementing 108N with acetic acid (AcOH)
Egg-to-pupa time
AB AB A B
Median time (dAEL)
Egg-to-pupa viability
A A A A
Survivorship (%)
108N
108N (0.45% AcOH)
108N (0.6% AcOH)
108N (0.75% AcOH)

**Supplementary Figure 3. Optimizing the 108N diet for larval development.** **a, c, e, g,** and **i,** Median egg-to-pupa duration in days after-egg-lay (dAEL) and mean percentage egg-to-pupa survivorship of flies reared on 108N supplemented with (**a**) 1 mg/L; 10 mg/L; or 100 mg/L carnitine (C), (**c**) 0.05%; 0.1%; or 0.5% coconut oil (CO), (**e**) 0.3 g/L; 0.1 g/L; or 0.03 g/L linolenic acid (LA), (**g**) 10%; 25%; or 50% sucrose (S), and (**i**) 0.45%; 0.6%; or 0.75% acetic acid (AcOH). **b, d, f,** and **h,** Median egg-to-adult duration in days after-egg-lay (dAEL) and mean percentage egg-to-adult survivorship of flies reared on 108N supplemented with (**b**) 1 mg/L; 10 mg/L; or 100 mg/L carnitine (C), (**d**) 0.05%; 0.1%; or 0.5% coconut oil (CO), (**f**) 0.3 g/L; 0.1 g/L; or 0.03 g/L linolenic acid (LA), and (**h**) 10%; 25%; or 50% sucrose (S). One-way ANOVA followed by Tukey's HSD test. Different letters represent statistically significant differences ($p < 0.05$).
